# “*mazEF* mediated programmed cell death in Escherichia coli”: is it?

**DOI:** 10.1101/048728

**Authors:** Bhaskar Chandra Mohan Ramisetty, Swati Raj, Dimpy Ghosh

## Abstract

Toxin-antitoxins systems (TAS) are prokaryotic operons containing two small overlapping genes which encode two components referred to as Toxin and Antitoxin. Involvement of TAS in bacterial programmed cell death (PCD) is highly controversial. MazEF, a typical type II TAS, is particularly implicated in mediating PCD in *Escherichia coli*. Hence, we compared the metabolic fitness and stress tolerance of *E. coli* strains (MC4100 and its *mazEF*- derivative) which were extensively used by proponents of *mazEF*- mediated PCD. We found that both the strains are deficient in *relA* gene and that the *ΔmazEF* strain has lower fitness and stress tolerance compared to wild type MC4100. Furthermore, these strains are likely not isogenic. We could not reproduce *mazEF* mediated PCD which emphasizes the need for skeptic approach to the PCD hypothesis.

## Introduction

Toxin-antitoxin systems (TAS) are autoregulated operons that consist of two small overlapping genes. One gene codes for a toxic protein, which impedes macromolecule synthesis, while the other gene codes for an RNA/protein that directly or indirectly alleviates the effects of the toxin (Gerdes and Maisonneuve 2012, Yamaguchi and Inouye 2011, Yamaguchi, *et al*. 2011). They are encoded by most prokaryotic genomes, often in multiple numbers (Anantharaman and Aravind 2003, Pandey and Gerdes 2005, Shao, *et al*. 2011). *mazEF* TAS, a typical Type II TAS, is particularly implicated in bacterial programmed cell death (PCD) (Aizenman, *et al*. 1996, Erental, *et al*. 2012, Kolodkin-Gal, *et al*. 2007). *mazEF* TA system is an operon consisting of two open reading frames (ORFs), *mazE* and *mazF* located downstream of *relA* and upstream of *mazG* in *Escherichia coli*. MazE, the antitoxin, physically binds to MazF, alleviating the toxic effects of the latter (Zhang, *et al*. 2003a). MazF is an endoribonuclease that cleaves RNAs independent of the ribosome with specificity for ACA sites (Zhang, *et al*. 2003b). *mazEF* operon is autoregulated by the MazEF complex (Marianovsky, *et al*. 2001). Stresses like nutrient starvation cause depletion of MazE proteins, leaving MazF free to act on its target RNAs. MazE is a substrate of ClpAP and is rapidly degraded as opposed to degradation of MazF. Conditions that prevent the expression of *mazEF* loci, transcription as well as translation, render MazF free of MazE. Several translational inhibitors also activate MazF (Sat, *et al*. 2001) culminating in the death of the cell. The dramatic decrease in the population determined by *mazEF* loci is interpreted as *mazEF*-mediated PCD. Engelberg-Kulka’s lab has pioneered the complex pathways of PCD involving factors like ppGpp (Aizenman, *et al*. 1996), Extracellular Death Factor (EDF) (Belitsky, *et al*. 2011, Kolodkin-Gal, *et al*. 2007) and Reactive Oxygen Species. The cleavage of 43 nucleotides from 3’-terminus of the 16S rRNA and also 5’-untranslated regions (UTRs) of specific transcripts by MazF allows bypassing the requirement of Shine-Dalgarno sequence for translational initiation (Vesper, *et al*. 2011). This allows the formation of “stress translation machinery” (Moll and Engelberg-Kulka 2012) which produces two sets of proteins, “survival proteins” and “death proteins” (Amitai, *et al*. 2009). The expression of death proteins, like SlyD, ClpP, YjiD, in addition to MazF mediated RNA degradation result in cell death observed as a dramatic decline in the colony forming units. *mazEF*-mediated PCD is induced by a plethora of stresses like antibiotics, heat shock and nutrient starvation (Amitai, *et al*. 2004, Sat, *et al*. 2001). Furthermore, *mazEF*-mediated PCD is an alternate pathway that inhibits “apoptosis like cell death” (ALD) in bacteria (Erental, *et al*. 2012). ALD is regulated by RecA-LexA pathway and hence is a part of SOS response. Recently, it was reported that RecA mediated autoproteolysis of LexA is modulated through the *mazEF*-mediated pathway (Erental, *et al*. 2014). It was theorized that the PCD phenomenon is a manifestation of “nutritional altruism”, meaning that the majority of the cells die to provide nutrition for surviving small subset of cells (Engelberg-Kulka, *et al*. 2006). It was also proposed that this kind of PCD could pave channels for the supply of nutrients during the formation of biofilms (Kolodkin-Gal, *et al*. 2009).

However, PCD phenomenon was irreproducible and it was also found that several inadvertent mutations were present around *mazEF* loci in the strains used to study *mazEF*-mediated PCD (Dorsey-Oresto, *et al*. 2013, Tsilibaris, *et al*. 2007). Several contradictions with regard to the protease that degrades MazE, the number of promoters in *mazEF* operon and the role of ppGpp were also reviewed (Ramisetty, *et al*. 2015). TAS are also implicated in a contrasting phenomenon called “persistence”, a form of non-inheritable tolerance to antibiotic stress exhibited by a small fraction of cells of a population (Moyed and Bertrand 1983). The increasing deletions of single TAS in *E. coli* were correlated with decreasing persistence, indicating that each TAS could confer a limited degree of persistence (Maisonneuve, *et al*. 2013, Maisonneuve, *et al*. 2011). MazF was shown to induce stasis and further enhance the formation of persisters (Tripathi, *et al*. 2014). TAS are proposed as attractive targets for novel antibacterial drugs. However, the fundamental objective for such drug designing still remains ill-defined, to activate or inactivate TAS. Hence, it is an important and immediate necessity to arrive at irrefutable conclusions regarding the role of *mazEF* TA system in PCD. In this study, we have independently analyzed the metabolic fitness and stress tolerance of MC4100 *relA*^+^ and *ΔmazEF relA*^+^ strains (Engelberg-Kulka, *et al*. 1998) that were extensively used to elucidate *mazEF*-mediated PCD.

## Materials and methods

### Strains and chemicals

*E. coli* MC4100*relA*^+^, *ΔmazEFrelA*^+^ strains (Engelberg-Kulka, *et al*. 1998) and MG1655 were used in this study. All chemicals were purchased from Himedia^TM^.

### SMG phenotype

16 hour old colonies of MC4100, *ΔmazEF* and MG1655 (control) strains, grown on Luria Bertani Agar (LA), were streaked on minimal A agar plates supplemented with and without a mixture of Serine, Methionine and Glycine (10 μg/mL each). Plates were incubated at 37°C for 24 hours.

### Growth curve

15 hour old overnight cultures were diluted 100 fold in LB medium and 2 μl of the diluted cultures were inoculated into 200 μl of LB in microtitre plate wells in triplicates. The microtitre plates were incubated at 37°C with 170 rpm shaking. Optical density at 595 nm was measured in a microtitre plate reader (Biorad^TM^), every 30 minutes, for 7 hours. Growth rate as a measure of change in OD per hour was calculated for each time point and plotted as a graph to infer the maximum growth rate attained by each of the strains.

### Prolonged stationary phase assay

3 μl overnight cultures of the strains were inoculated into tubes containing 3 mL of LB broth. The tubes were incubated at 37°C with 170 rpm shaking. Samples were taken at an interval of 24 hours for 6 days and the cells were plated after appropriate dilutions and colonies were counted. The growth of MC4100 at 24 hours was taken as 100% (maximum CFU of all the strains). All other readings of strains at different time points were percent CFU relative to CFU of MC4100 in 24 hours sample.

### Biofilm assay

Overnight cultures were diluted 100 fold in fresh LB tubes. 200 μl of LB broth was put in 96 well microtitre plate and inoculated with 2 μl of each strain. The plates were incubated at 37°C for 16, 24, 48 and 72 hours at 37°C. After the specified time points the plates were washed with PBS to remove floating cells. 125μl of 1% crystal violet was added to each well and left for 20 minutes. The plates were washed with water twice and the dye was re-dissolved by adding 90% ethanol and reading were taken at 595nm and readings were represented as the amount of biofilm formation. Experiments were carried out independently thrice in quadruplicates.

### Osmotic stress

Overnight cultures were diluted 1000 fold in fresh LB tubes containing 1% (control), 2% and 3% NaCl. They were allowed to grow for 24 hours and were serially diluted with 0.9% NaCl. 100μl of diluted culture was spread on LA plates. Colonies were counted and CFU was calculated according to the dilutions made. Maximum CFU in the given amount of media (obtained for MC4100 at 24 hours) was taken as 100% and that of other strains/conditions as relative percentages. For the sensitivity of strains on agar plates, several LB agar plates were made with increasing amounts of NaCl concentrations (1% control, 2% and 3%). Colonies were streaked on these plates and incubated for 24 hours at 37°C.

### PCD assay

Overnight cultures of strains were inoculated into fresh 100 mL LB broth (1000 fold dilution) and grown in LB medium at 37°C with 170 rpm in flasks till mid log phase was reached. 3 mL of each culture was distributed into four tubes and to induce stress, 16 μg/mL of Chloramphenicol or 15 μg/mL of nalidixic acid were added or the tubes were incubated at 50°C for 60 min in a water bath without shaking. Samples were appropriately diluted and plated on LA and incubated at 37°C. Colonies were counted and presented as percent CFU relative to CFU of MC4100 of the control.

### Antibiotic sensitivity test

Conventional disc diffusion method was used to measure relative sensitivity of the strains. 100μl of diluted (100 fold) overnight cultures was spread on LB agar (height – 5mm) contained in plates with diameter 9.5cm. Premade antibiotic discs with defined concentrations (purchased from HiMedia^TM^) were placed on the agar plates after 20 minutes. The plates were incubated overnight at 37°C. Diameters of the zones of inhibition were measured and the graph was plotted.

## Results and Discussion

### SMG phenotype and growth rates of MC4100 and *ΔmazEF* strains

ppGpp, the alarmone produced by RelA, induces stringent response to mitigate nutrient starvation primarily by reducing the synthesis of macromolecules. However, *E. coli* MC4100 strain harbours a *relA1* (Metzger, *et al*. 1989) mutation that impairs the production of ppGpp and is phenotypically characterized by an inability to grow on minimal media plates supplemented with Serine, Methionine and Glycine (SMG phenotyping) (Uzan and Danchin 1976). ppGpp was shown to be essential to initiate *mazEF*-mediated PCD in *E. coli* MC4100 strain and hence the proponents of “*mazEF*-mediated PCD” constructed MC4100 *relA*^+^ and *ΔmazEFrelA*^+^ (Aizenman, *et al*. 1996, Engelberg-Kulka, *et al*. 1998, Hazan, *et al*. 2004). However, both these strains were reported to be *relA* mutants (Tsilibaris, *et al*. 2007). Therefore, we revaluated the strains for *relA*^+^ using SMG phenotyping. We found that both the strains were SMG negative (Fig 1A), implying that the curing of *relA1* mutation to *relA*^+^ was not successful, corroborating the findings of Tsilibaris *et al.* 2007. Unfortunately, the construction of *relA*^+^ derivatives of MC4100 and its *ΔmazEF* strains were not detailed (Engelberg-Kulka, *et al*. 1998). It should be noted that the conversion of *ΔmazEFrelA1* to *ΔmazEFrelA*^+^ using lysate grown on any *E. coli* (*mazEF^+^relA*^+^) strain is highly improbable using phage mediated generalized transduction. This is because *relA* and *mazEF* ORFs are separated by 77 nucleotides, meaning that they are highly linked. In the following sections, these strains are called as MC4100 and *ΔmazEF*.

**Fig 1A.**
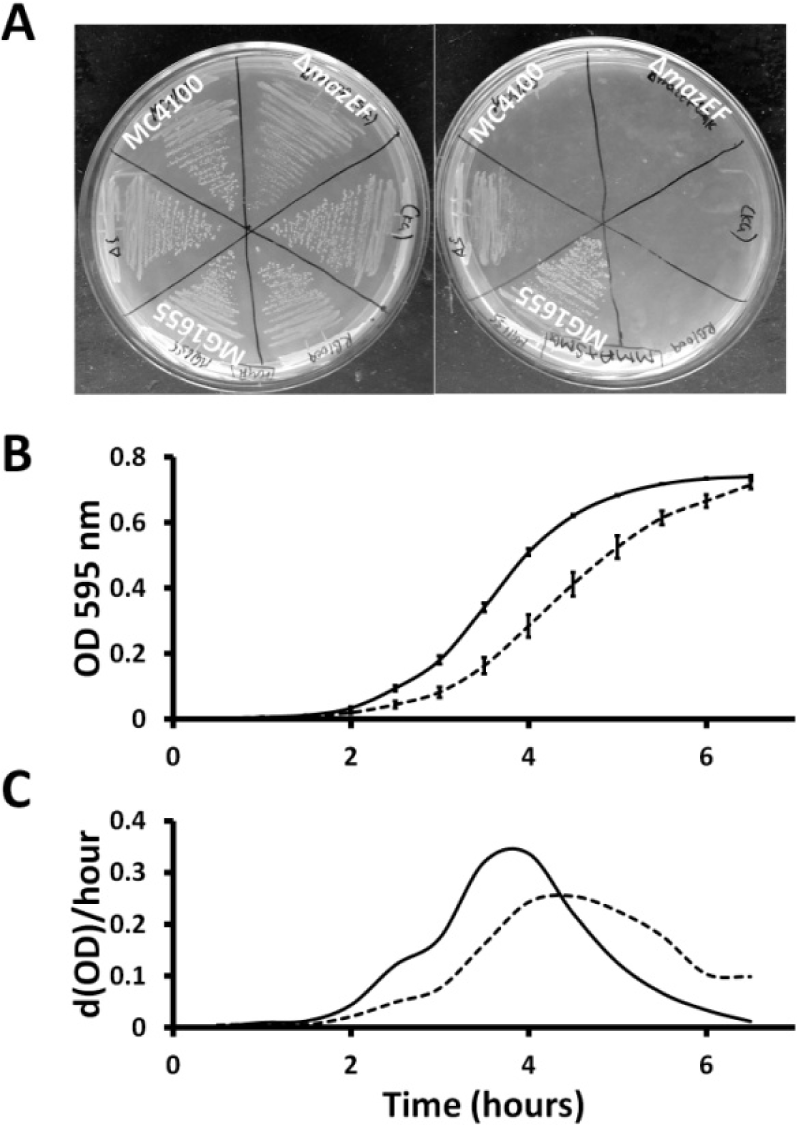
SMG phenotyping of MC4100*relA*^+^ and *ΔmazEFrel*^*A*+^. Plate on the left is minimal A media and plate on the right is minimal A media supplemented with serine, methoinine and glycine (10μg/ml each), **B**. Growth curve of MC4100*relA*^+^ (solid line) and *ΔmazEFrelA*^+^ (dotted line) in LB medium at 37°C with 170 rpm. The result presented is a representative of one experiment, out of three, done in triplicates in a microtitre plate. **C**. The growth rate of MC4100*relA*^+^ (solid line) and *ΔmazEFrelA*^+^ (dotted line) calculated as the change in OD per hour at each time point for the growth curve in B.

The growth rate of a bacterial strain in optimal conditions is one fundamental indicator of metabolic fitness. Hence, we determined the growth rate of MC4100 and *ΔmazEF* strains in optimal conditions. We found that the maximum growth rates attained by MC4100 and *ΔmazEF* were 0.35 and 0.25 OD/hour respectively (Fig 1B, C). This 29% lower maximal growth rate of *ΔmazEF* strain compared to MC4100 indicates that the *ΔmazEF* strain has decreased fitness. Theoretically, during optimal conditions the phenotypic expression of TA loci is expected to be minimal and hence should not hinder the growth rate. In contrast to these expectations, we found that the *ΔmazEF* strain had lower growth rate compared to the WT implicating the known genetic manipulations done (Aizenman, *et al*. 1996, Engelberg-Kulka, *et al*. 1998) at this loci. Furthermore, the colonies of *ΔmazEF* strain are consistently smaller than the wild type (data not shown).

### Analysis of MC4100 and *ΔmazEF* strains in prolonged stationary phase and biofilm formation

The ability of a bacterial strain to grow to maximum colony forming units (CFU) in a given volume of nutrients, under optimal conditions, is an indication of its metabolic ability to use the resources efficiently and also sustain during nutrient exhaustion. We performed long term survival assay of both the strains for a period of 6 days. In 24 hours MC4100 yielded 1.74 x10^12^ CFU/ml (normalized as 100%), while *ΔmazEF* yielded 8.7 x 10^11^ CFU/ml (50% of WT) (Fig 2A). Over the period of 6 days, CFU of both the strains declined several folds. We found that *ΔmazEF* strain had 30-50% lower CFU compared to similarly growing WT during the first five days and equalled CFU of WT on the 6^th^ day. CFU of *ΔmazEF* has remained relatively lesser in our prolonged stationary phase experiments. This indicates that *ΔmazEF* strain is less efficient in metabolizing the nutrient resources.

**Fig 2A.**
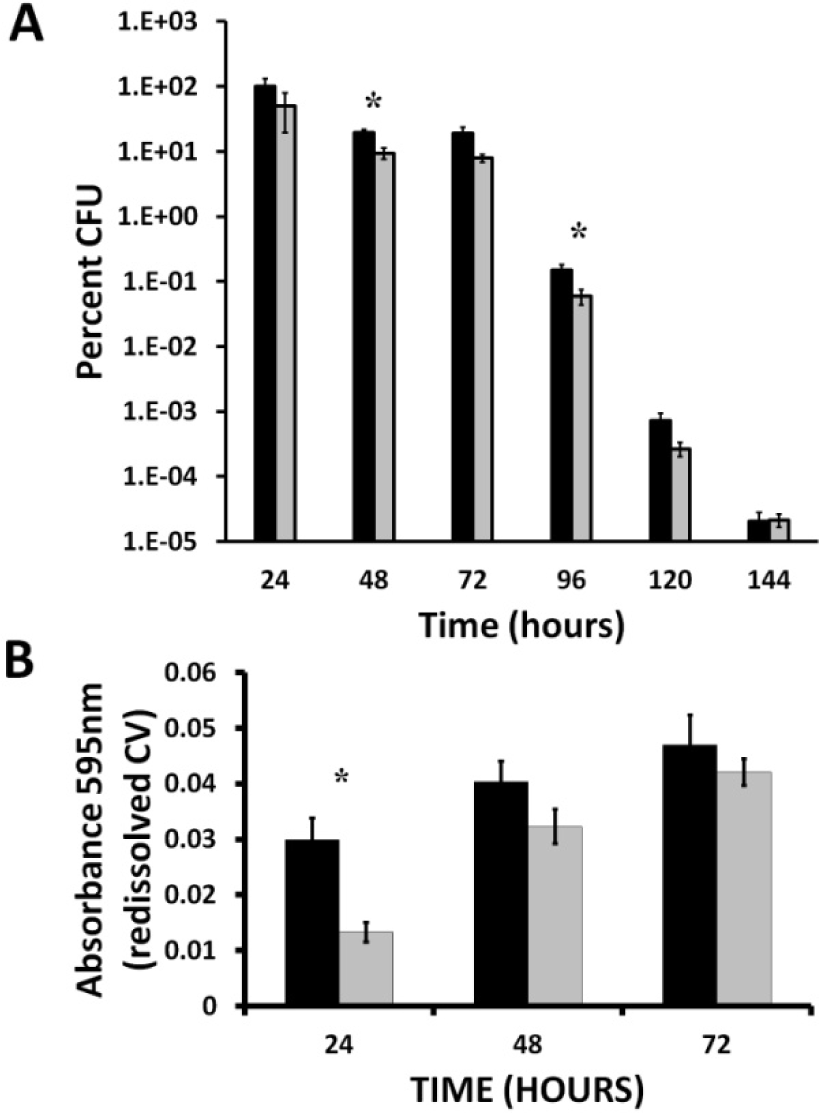
Total viable counts of MC4100*relA*^+^ (solid bars) and *ΔmazEFrelA*^+^ (grey bars) grown at 37°C with 170 rpm for 6 days presented as percent CFU on a log scale. *Two-tailed P value = 0.0089 and 0.0495 for 48 and 96 hours respectively. **B**. Biofilm formation of MC4100*relA*^+^ (solid bars) and *ΔmazEFrelA*^+^ (grey bars) for 24, 48 and 72 hours of incubation at 37°C in microtitre plates. Both the bar diagrams are one representative experiment done in quadrulicates. Error bars indicate standard errors. *Two-tailed P value = 0.0019.

Biofilm formation is a feature exhibited by most free living bacteria when grown in relatively stagnant liquid habitats. The ability to form persistent biofilms is an ecological strategy to withstand stresses like antibiotics (Hoiby, *et al*. 2011, Lewis 2008). Several TAS enhance the formation of biofilms (Norton and Mulvey 2012, Van Acker, *et al*. 2014, Wang and Wood 2011). It was observed that the MC4100 exhibited enhanced biofilm formation but was composed largely of dead cells compared to *ΔmazEF* strain (Kolodkin-Gal, *et al*. 2009). It was reported that *mazEF* and *dinJ-yafQ* TAS are involved in cell death during formation of biofilms. We performed similar biofilm assays for a prolonged duration (24, 48 and 72 hours) and found that the biofilm formation increased with incubation time for MC4100 and *ΔmazEF* strains. We observed that *ΔmazEF* strain formed 50% less biofilm compared to that of WT after 24 hours. We also noted that biofilm formation of both the strains increased at similar rates. If the majority of the cells in the WT biofilm were dead as reported by Kolodkin-Gal *et al.*, 2009, one would expect that MC4100 does not form anymore biofilm upon prolonged incubation. Our results indicate that biofilm formed by MC4100 may be composed of proportionate viable cells.

### Stress tolerance of MC4100 and *ΔmazEF* strains

Even though numerous TAS are characterized, so far *mazEF* system is the only TAS shown to mediate PCD in *E. coli*. However, a study (Tsilibaris, *et al*. 2007) failed to reproduce this PCD phenomenon using the same strains used by Engelberg-Kulka’s lab (Engelberg-Kulka, *et al*. 1998). Hence, we performed PCD assay to independently test the hypothesis with the same strains from Engelberg Kulka’s lab. Different samples from exponentially growing cells were treated with Chloramphenicol (16 μg/mL), Nalidixic acid (15 μg/mL) or were subjected to 50°C heat shock for one hour. Percent CFU was normalized to MC4100 untreated sample and presented on a log scale (Fig 3A). In contradiction to the studies that have established *mazEF*-mediated PCD (Engelberg-Kulka, *et al*. 2004), *ΔmazEF* strain was 36-54% more sensitive relative to similarly treated WT, in all the conditions tested. We do not have an explanation for this contradiction other than that some unobvious critical factor in the experimental setup may not have been followed by us or occurrence of any spontaneous mutations in the strains used during transport and storage. However, our findings corroborate observations reported by Laurance Van Melderen’s lab (Tsilibaris, *et al*. 2007).

**Fig 3A.**
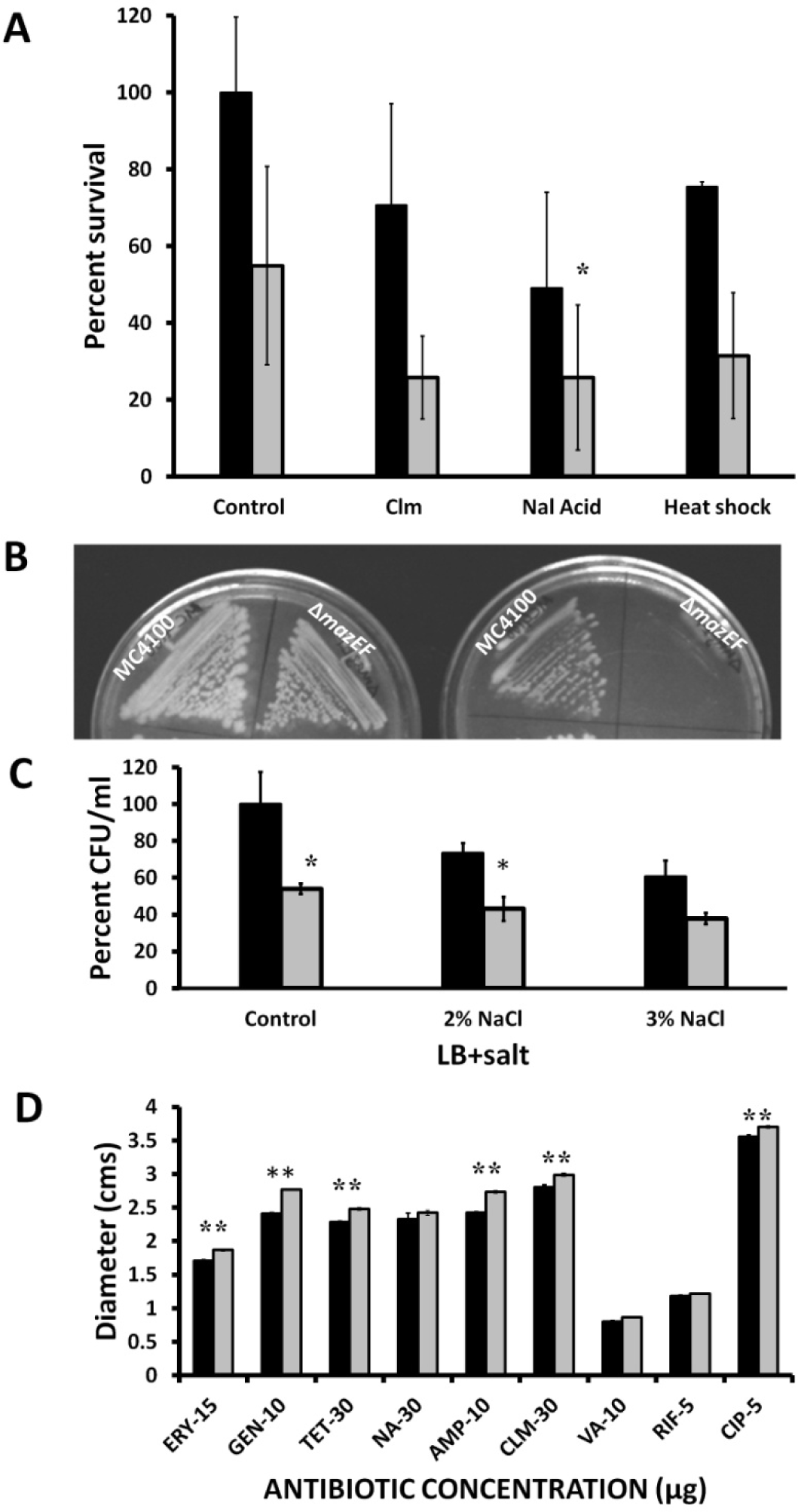
Stress tolerance of the MC4100*relA*^+^ (solid bars) and *ΔmazEFrelA*^+^ (grey bars) to various stresses. Midlog cultures were distributed into fresh tubes and exposed to different stresses (clm = chloromphenicol 16 μg/ml, NA = Nalidixic acid 15 μg/ml, Heat shock at 50°C) each for one hour. Samples were taken and plated after dilution, colonies counted and presented as percent survival relative to MC4100 in control. The graph is an average of three independent experiments and the error bars indicate standard error. *Two tailed P value equals 0.0168. **B**. The sensitivity of MC4100*relA*^+^ (solid bars) and *ΔmazEFrelA*^+^ (grey bars) to antibiotics by disc diffusion method. Overnight cultures were spread on LB agar plates and antibiotic discs were placed on the plates and incubated at 37°C for 24 hours. Diameters of the zones of inhibition around each disc were measured at different angles. The shown graph is an average of three independent experiments. Ery = Erythromycin (15 μg), Gen = Gentamycin (10 μg), Tet = Tetracycline (30 μg), NA = Naldixic acid (30 μg), Amp = Ampicillin (10 μg), Clm + Chloramphenicol (30 μg), VA = Vancomycin (10 μg), Cip = Ciprofloxacin (5 μg). ** Unpaired two-tailed P value < 0.0001 in all significant cases.

To determine the tolerance of these strains to osmotic stress we planted them on LA plates with increasing amounts of NaCl (1%, 2% and 3%). As shown in Fig 3B, the colonies became increasingly water-filled with increasing salt concentration. Interestingly, we found that *ΔmazEF* strain showed relatively poor growth on LB agar plates containing 3% NaCl. Similarly, when grown in aerated liquid cultures with 1%, 2%, 3% NaCl concentrations for 24 hours at 37°C, *ΔmazEF* had 50-60% lower CFU, relative to WT, in all NaCl concentrations tested (Fig 3C). Contrary to the expectation that both strains could be sensitive to NaCl due to *mazG* deletion, we found that *ΔmazEF* strain is relatively sensitive to osmotic stress. This experiment does show that *ΔmazEF* strain may have an additional mutation, spontaneous and/or inadvertent mutation. It was shown that *mazEF* mediates PCD upon encountering antibiotics (Amitai, *et al*. 2004, Hazan, *et al*. 2004, Kolodkin-Gal and Engelberg-Kulka 2006). Recent studies implicate TAS in persistence under antibiotics stress (Maisonneuve, *et al*. 2011) and a study (Tripathi, *et al*. 2014) specifically implicates *mazEF* in the formation of persisters. Hence, we analyzed the sensitivities of MC4100 and *ΔmazEF* strains to various antibiotics. We chose the simple method of disc diffusion to test the relative sensitivity of the strains for analysis. We observed that *ΔmazEF* strain was relatively sensitive to all the antibiotics tested except for Rifampicin and Vancomycin (negligible zone of inhibition) (Fig 3D). Contrary to our anticipation that MC4100 would be more sensitive based on PCD hypothesis, we found that *ΔmazEF* strain was relatively sensitive to antibiotics.

Our study emphasizes that *ΔmazEF* strain is physiologically deficient relative to its wild type MC4100. “*mazEF*-mediated PCD” is linked with oxidative stress (Hazan, *et al*. 2004) and DNA repair (Erental, *et al*. 2014, Erental, *et al*. 2012) and so are MazG homologues (Lu, *et al*. 2010, Lyu, *et al*. 2013). MC4100 and MC4100*ΔmazEF* strains (Engelberg-Kulka, *et al*. 1998), used to demonstrate “*mazEF*-mediated PCD” by Engelberg Kulka’s lab, were reported to have a deletion in *mazG* ORF as well (Gross, *et al*. 2006, Tsilibaris, *et al*. 2007). *mazG*, a highly conserved gene, encodes a nucleoside triphosphate pyrophosphohydrolase which hydrolyzes NTPs and dNTPS to their monophosphates and inorganic pyrophosphates (Zhang and Inouye 2002). Recently it was found that Mycobacterial MazG “safeguards” the genome by degrading 5-OH-dCTP, thereby preventing CG to TA mutation (Lyu, *et al*. 2013). Several MazG homologues are implicated in stresses like oxidative stress (Lu, *et al*. 2010) and salt stress (Culligan, *et al*. 2012). Hence, it is imperative to address the deletion of the *mazG* gene in both the strains and important to experimentally demonstrate the proposed nutritional altruism as a consequence of *mazEF*-mediated PCD. We conclude that the strains, MC4100 and *ΔmazEF* strain used in PCD studies are not isogenic. PCD phenomenon is not reproducible in our hands and may need further scrutiny. Probably the whole genome sequencing of these strains would reveal the explanations for several observations. It is also suggest to refrain from using these strains (MC4100 and *ΔmazEF* (Engelberg-Kulka, *et al*. 1998)) in genetic and physiological studies until a complete genotype of these strains is established.

## Acknowledgment

We thank Prof. Hanna Engelberg Kulka for the kind donation of the strains. We thank the management of SASTRA University for providing Prof T. R. Rajagopalan grants and Central Research Facility.

